# Evidence for the role of transcription factors in the co-transcriptional regulation of intron retention

**DOI:** 10.1101/2021.11.18.469150

**Authors:** Fahad Ullah, Maayan Salton, Anireddy SN Reddy, Asa Ben-Hur

## Abstract

Alternative splicing is a widespread regulatory phenomenon that enables a single gene to produce multiple transcripts. Among the different types of alternative splicing, intron retention is one of the least explored despite its high prevalence in both plants and animals. The recent discovery that the majority of splicing is co-transcriptional has led to the finding that chromatin state affects alternative splicing. Therefore it is plausible that transcription factors can regulate splicing outcomes. We provide evidence for this hypothesis by studying regions of open chromatin in retained and excised introns. Using deep learning models designed to distinguish between regions of open chromatin in retained introns and non-retained introns, we identified motifs enriched in IR events with significant hits to known human transcription factors. Our model predicts that the majority of transcription factors that affect intron retention come from the zinc finger family. We demonstrate the validity of these predictions using ChIP-seq data for multiple zinc finger transcription factors and find strong over-representation for their peaks in intron retention events.

**Availability:** Source code available at https://github.com/fahadahaf/chromir

## 1. Introduction

Alternative splicing is a widespread regulated phenomenon that enables a single gene to encode structurally and functionally different transcripts [1,2]. The primary forms of alternative splicing are exon skipping, intron retention (IR), and alternative 3′ and 5′ splicing. While exon skipping is well studied, intron retention remains an under-appreciated phenomenon [3]. IR is the primary form of alternative splicing in plants [4,5], and recent studies have shown it to have a high prevalence in human [6, 7]. Many disease-causing mutations are pathogenic through their effect on splicing, often leading to IR [6,8,9]. For example, IR is associated with genetic variants with deleterious effect on the function of tumor suppressor genes [10].

In recent years, efforts have been made to understand the regulation of IR and the factors that contribute to it. Braunschweig et al. [7] recently published a draft “IR splicing code”: a predictive model that uses a total of 136 features thought to be associated with IR in mammals. These features include base composition of an intron and its flanking exons, features that describe gene architecture, and splice site strength. This model is limited in that it ignores sequence elements that contribute to the regulation of IR. The discovery that splicing occurs co-transcriptionally suggests that chromatin state might be relevant to alternative splicing [11,12]. Recent work provides evidence for the regulatory contribution of chromatin state to exon skipping [13], and our labs have provided preliminary evidence for its role in regulating IR in plants [14]. Open chromatin is one of the most important signatures for the study of chromatin structure. One of the primary tools for probing open chromatin is through exposure of DNA to deoxyribonuclease I (DNase I), which is an enzyme that cleaves DNA. Regions of the genome that are sensitive to its action—DNase I hypersensitive sites (DHSs)—have been used as an indicator of chromatin accessibility *in-vivo* [15]. DHSs have been used extensively to identify several types of regulatory elements such as promoters, enhancers, silencers, and insulators [16,17]. Furthermore, when a regulatory protein binds DNA, it protects it against the action of DNase I [18] and leaves a footprint which can be identified using DNase I-seq data [19,20]. When it comes to alternative splicing, Mercer et al. [13] have shown an association between DHSs and exon-skipping, reporting that higher numbers of DHS-containing exons are alternatively spliced. Furthermore, this study reports that DHS exons with promoter and enhancer-like features have a higher fractional overlap with alternative splicing. Braunschweig et al. [7] explored the co-transcriptional regulation of splicing, reporting higher chromatin accessibility in retained introns and that polymerase II elongation speed affects IR and vice-versa. In another work, it has been reported that zinc finger transcription factors (TFs) have a regulatory role in exon skipping [21]. Recently, we studied the association between chromatin accessibility and intron retention in plants [14]. We identified potential regulatory elements occurring primarily in the 3’ flanking exons of IR events, several of which significantly match plant zinc finger binding site motifs. As further motivation for considering the role of TFs in splicing regulation, we explored the frequency of motif matches for different TF families across regions of open chromatin in the human genome. We observe that the prevalence of motif matches in human intragenic regions is comparable in number to the overall number of matches in intergenic regions, even without controlling for their much greater length (see Figure 1); a similar observation was made in plants [22]. This suggests a regulatory role of TFs beyond the regulation of gene expression.

**Figure 1.**
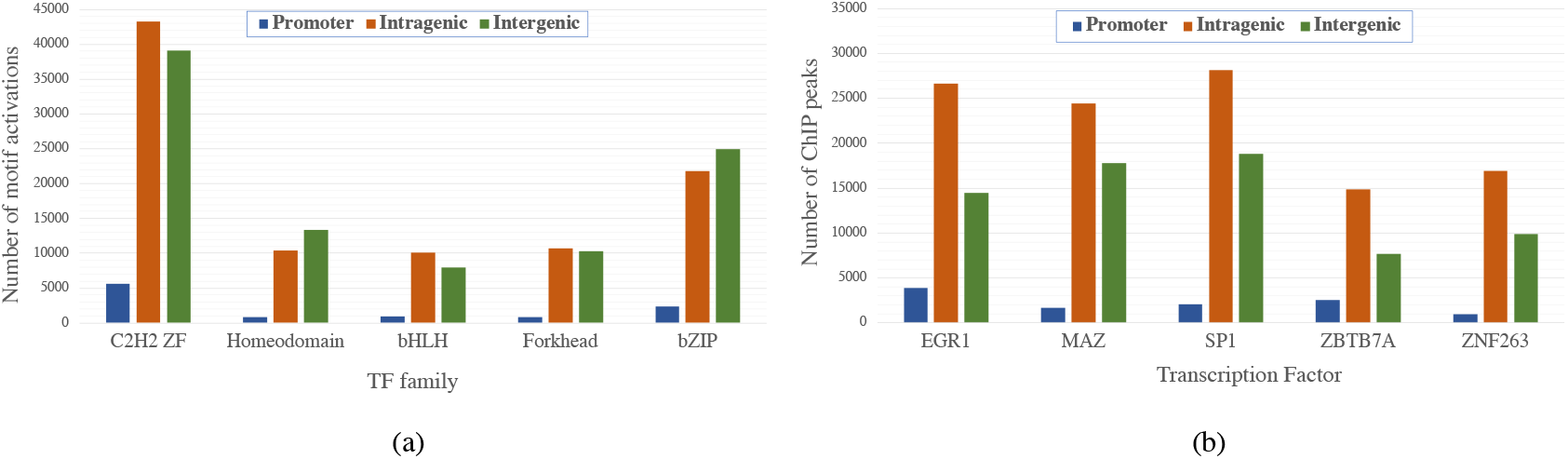
TF binding sites across the genome. (a) The number of predicted binding sites for different TF families in the promoter, intragenic, and intergenic regions of the human genome. These counts were obtained by training the Basset-like network [23] and analyzing the motifs learned by the network (see supplementary methods for more details). (b) ChIP peak counts for five human zinc finger TFs in the promoter, intragenic, and intergenic regions of the human genome.

Deep neural networks have become the tool of choice for exploring complex phenomena such as chromatin accessibility its regulatory roles [23–26]. A remarkable advantage of these models is their ability to capture the underlying patterns in large noisy datasets directly from sequence with minimal pre-processing, learning motifs of the regulatory proteins involved as part of the training process. Deep learning has been used in genomics for TF binding prediction [27–29], chromatin accessibility analysis [23–25], prediction of chromatin structure and its modifications [30,31], identification of RNA-binding protein sites [32,33], and alternative splicing [34–36].

In this study we demonstrate that deep learning models can distinguish with good accuracy regions of open chromatin associated with IR from other intragenic regions of open chromatin. By analyzing the motifs learned by the network, we find that specific families of TFs are associated with IR events, mostly members of the zinc finger family of TFs; results of ChIP-seq experiments for multiple zinc finger TFs in the K562 cell line, one of three tier 1 ENCODE cell lines, support our findings for this association. Our work provides convincing evidence for a novel role of TFs in the regulation of IR, proposing a direction for further research.

## 2. Results

### 2.1. DHSs associated with IR can be accurately predicted from their sequences

In order to discover the sequence elements that regulate IR via its coupling with chromatin state we trained and evaluated deep learning models to distinguish DHSs associated with IR from non-IR DHSs in human and assessed and compared their performance. IR DHSs are regions in which a DHS overlapping an IR event was detected in at least one DNase I-seq experiment in a compendium of 164 samples described in the Methods section. Non-IR DHSs are intronic regions exhibiting a DHS where no IR is known to occur. Our primary focus is the purely convolutional architecture shown in Figure 2, that has demonstrated its effectiveness for predicting chromatin accessibility by Kelley et al. [23]. The model hyperparameters were tuned for our problem as described in the Methods section. Using this model we obtained accuracy of 0.54 as measured using the area under the precision-recall curve (AUC-PRC) (see Figure 3(a)). A more sophisticated model that uses a combination of convolutional and recurrent layers with multi-head attention achieved the same level of accuracy (see Supplementary Figure F2). We note that both deep-learning architectures outperformed a baseline approach that uses the gkm-SVM method [37]. This method achieved an AUC-PRC of 0.50. For additional results, including ROC curves, see Supplementary Figure F2.

**Figure 2.**
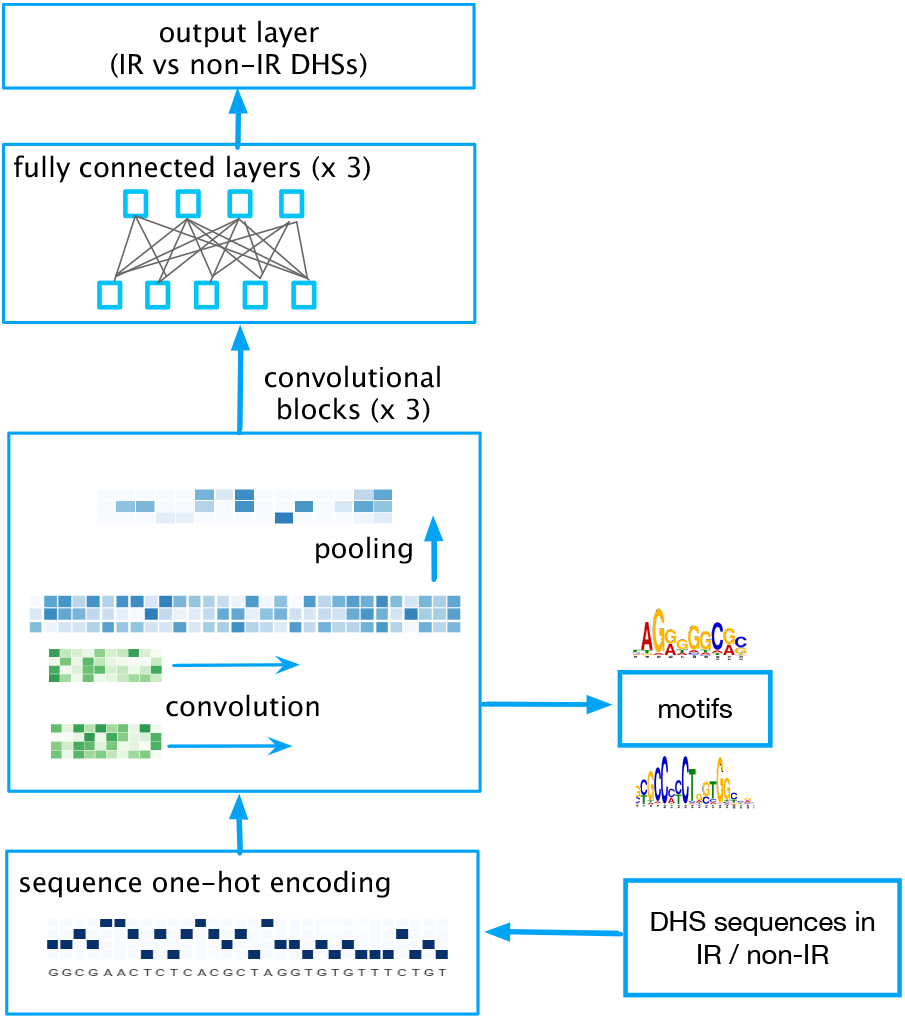
A deep learning model for predicting whether a region of open chromatin exhibits IR. The model receives as input the sequences of intragenic DHSs labeled as associated with IR or non-IR; the one hot encoding is processed through three layers of convolution, followed by three fully connected layers and the output layer that predicts a binary response that indicates whether a DHS exhibits IR or not. The convolutional filters of the first layer are used to extract Position Weight Matrices (PWMs) that are searched against a database of known TFs.

**Figure 3.**
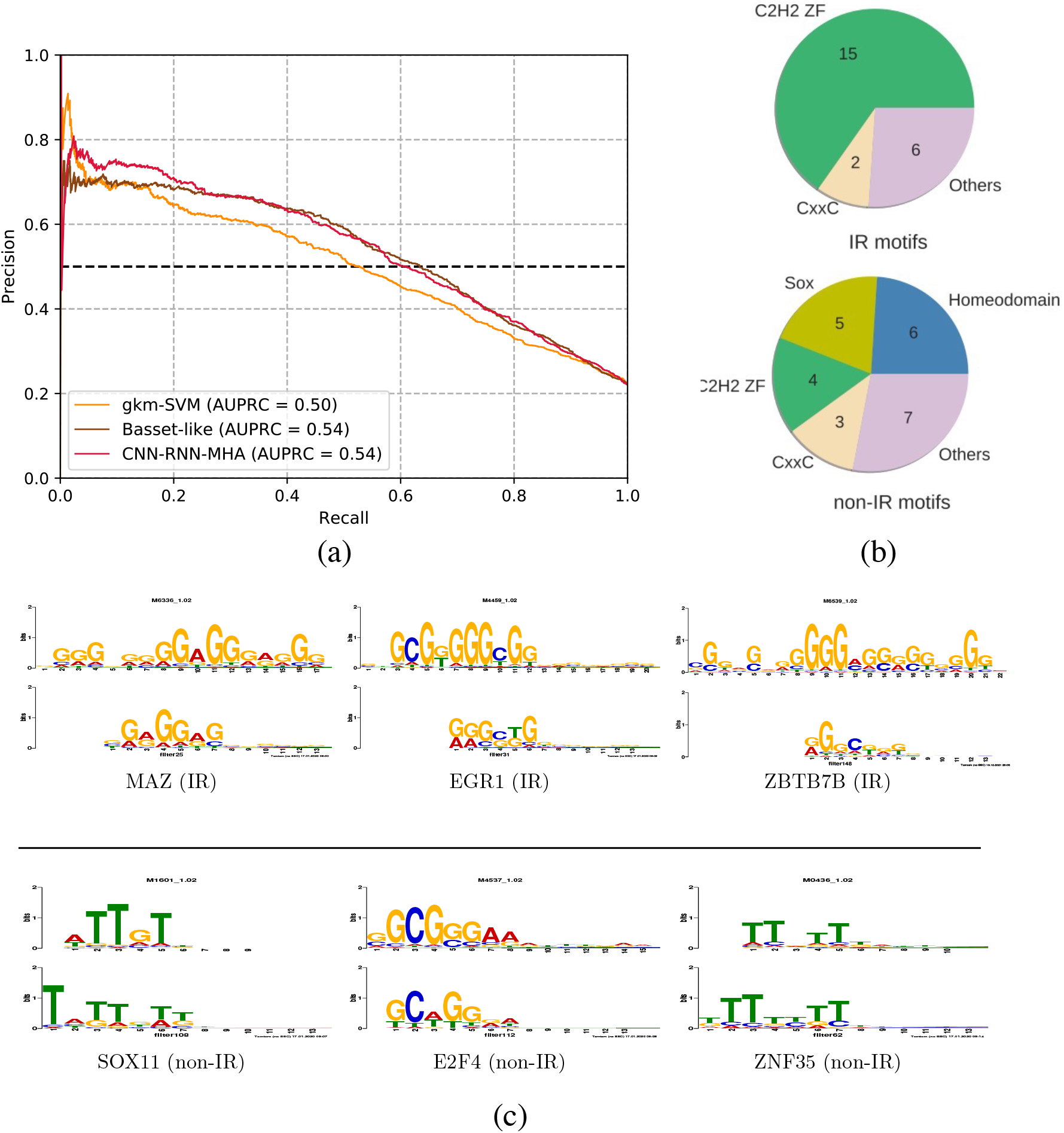
Classification accuracy and motifs detected by the network. (a) Precision-Recall curves for the two deep learning architectures and the gkm-SVM. The AUC-PRC values are also provided in the legend. (b) The distribution of TF families enriched in IR vs non-IR events. (c) The top three matches for the IR and non-IR convolutional layer filters against the CISBP database. In each match, the known TF motif is shown in the top row and the bottom row shows the CNN filter/motif. The motifs shown above the line are associated with IR DHSs, and those below the line are associated with non-IR DHSs.

Our results were generated using a one-hot encoding of the sequence of DHS regions. We note that word2vec embeddings provided a boost in accuracy, as shown in Supplementary Figure F2. However, this came at a cost of reduced interpretability of the models, leading to reduced ability to infer motifs associated with the learned convolutional filters. Therefore we chose to focus on models that used one-hot encoding as input.

### 2.2. The Zinc finger family of TFs are enriched in IR events

The filters of convolutional networks can be readily interpreted as motifs. To do so, we implemented the strategy described elsewhere [23,27] (see Methods section for more details). We analyzed the motifs that were derived from the convolutional filters for both the top positive and the top negative examples and searched both sets of motifs against the Human CIS-BP TF database [38] using the TomTom [39] tool. In case of IR DHSs, 23 motifs had significant hits against multiple known human TFs at a q-value < 0.01. In comparison, 25 of the non-IR motifs had significant matches. Figure 3(c) shows some of the top hits reported for both IR and non-IR motifs. We observe that most of the IR motifs have significant hits in the C2H2 zinc finger family of TFs (C2H2 ZF). Non-IR motifs on the other hand, are predominantly matched to the Homeodomain and Sox families of TFs (see Figure 3(b)). C2H2 ZF is the largest family of TFs and is highly active in the promoter, intergenic, and intragenic regions of the human genome (see Figure 1). However, it is highly significant to observe its enrichment in IR events compared to non-IR events. Zinc finger TFs have previously been implicated in the regulation of alternative splicing [21], particularly exon skipping. Here we report a role of this family in the regulation of intron retention. We note that some of our filters do not match a unique transcription factor. For example, the filter that matched MAZ, was also a good match for ZNF263. This is not surprising due to the similarity of the binding sites of zinc finger TFs. In what follows we provide additional evidence for the role of zinc finger TFs in regulating IR.

### 2.3. TF ChIP-seq analysis supports model predictions

To validate our findings using experimental data, we downloaded K562 ENCODE ChIP-seq datasets for all the zinc finger TFs identified by our model, resulting in six datasets. Using these datasets, we tested TF binding enrichment in IR vs. non-IR events, following a strategy similar to our previous work [14]: For each TF, we measured the overlap of its ChIP-seq peaks with IR and non-IR events and tested its significance using the Fisher-exact test. All the TFs demonstrated highly significant enrichment in IR events (see Table 1), validating our *in-silico* findings that the C2H2 ZF family plays a role in the regulation of IR.

**Table 1.** Enrichment of C2H2 ZF TF binding in IR vs non-IR events quantified using ChIP-seq peaks of the corresponding TF.

| TF | IR TF occupancy (%) | non-IR TF occupancy (%) | p-value |
| --- | --- | --- | --- |
| EGR1 | 12.51 | 7.16 | 1.27E-45 |
| MAZ | 11.42 | 6.06 | 9.75E-51 |
| ZBTB7A | 10.6 | 5.52 | 3.15E-49 |
| SP1 | 3.04 | 1.53 | 7.64E-16 |
| SP2 | 1.32 | 0.77 | 7.21E-06 |
| ZNF263 | 1.14 | 0.67 | 2.81E-05 |

To obtain additional insight, we plot a score that reflects the average occupancy of MAZ and EGR1 in retained introns and compare it with the patterns observed in non-retained introns using the same ENCODE ChIP-seq data used above, following a method described in the Methods section. MAZ and EGR1 were chosen as the TFs that exhibited the most significant level of enrichment in IR events. The resulting plot, shown in Figure 4, shows that for both TFs the occupancy is higher in the flanking exons of the retained introns. In contrast, in non-IR events, both EGR1 and MAZ are preferentially bound in the intronic regions. These TF-occupancy profiles suggest that these TFs have a role in both IR and non-IR events, but that their mechanism of action is different in each case. Further work is needed to test this hypothesis.

**Figure 4.**
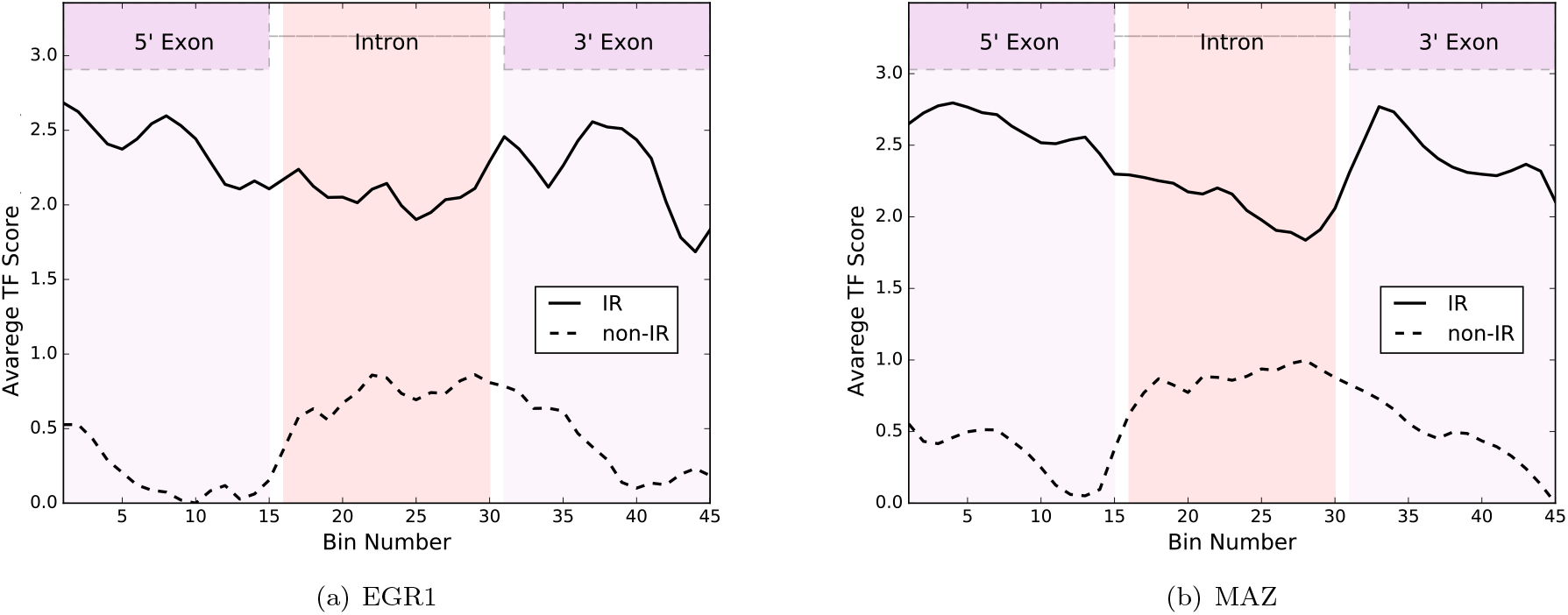
TF occupancy profiles across IR and non-IR events. These are shown for EGR1 (a) and MAZ (b). Binding location in each ChIP-seq peak was determined by the best match of the PWM of the corresponding TF, and the y-axis is the TF occupancy score as described in the Methods section.

### 2.4. Regulatory interactions between TFs in IR events

It is well known that TFs often function in tandem with each other to regulate their targets. To extract such regulatory interactions we have recently developed a method called SATORI to interpret deep architectures that use *attention* layers and extract statistically significant interactions between its convolutional filters [40]. SATORI uses the so-called attention matrix, which encodes relations between parts of the sequence; subsequent analysis of the convolutional filters that are active provides a profile of interactions between pairs of TFs that are associated with those filters. By comparing those profiles to those in a background set of sequences, we obtain interactions that are statistically significant. Using SATORI, with the negative examples as a background set to assess statistical significance we detected over 400 TF interactions in DHSs associated with IR at a significance level of 0.05. The top 20 predictions are shown in Figure 5, and the complete list is provided in supplementary table S2, A majority of these interactions involve the C2H2 ZF family, which is expected in view of C2H2 ZF TFs having the most hits from our model. To validate these interactions, we searched for matches in annotated interactions in the TRRUSTv2 [41] database that annotates TF regulatory roles and their interactions by text-mining the biomedical literature. Of the interactions detected by our model, we found 23 overlapping interactions in TRRUSTv2, which currently contains 8324 interactions. This is highly significant, with a p-value equal to 0 in a hypergeometric test. We also obtained significant overlap with protein-protein interactions from the HIPPIE database [42]: 17 of the detected interactions had support in HIPPIE, with a hypergeometric p-value < 1*e* − 52. The interactions overlapping with TRRUSTv2 and HIPPIE database are listed in Supplementary table S3 and S4, respectively. We also looked at the average distance between motifs predicted to interact and found that TF motifs preferentially interact in proximity, with a median distance of 120 bp, which is significantly less than what we would expect by chance (Mann-Whitney U test, p-value = 3.65*e* − 13). These results suggest that regulation of IR is orchestrated by complex interactions among TFs, predominantly from the C2H2 ZF family.

**Figure 5.**
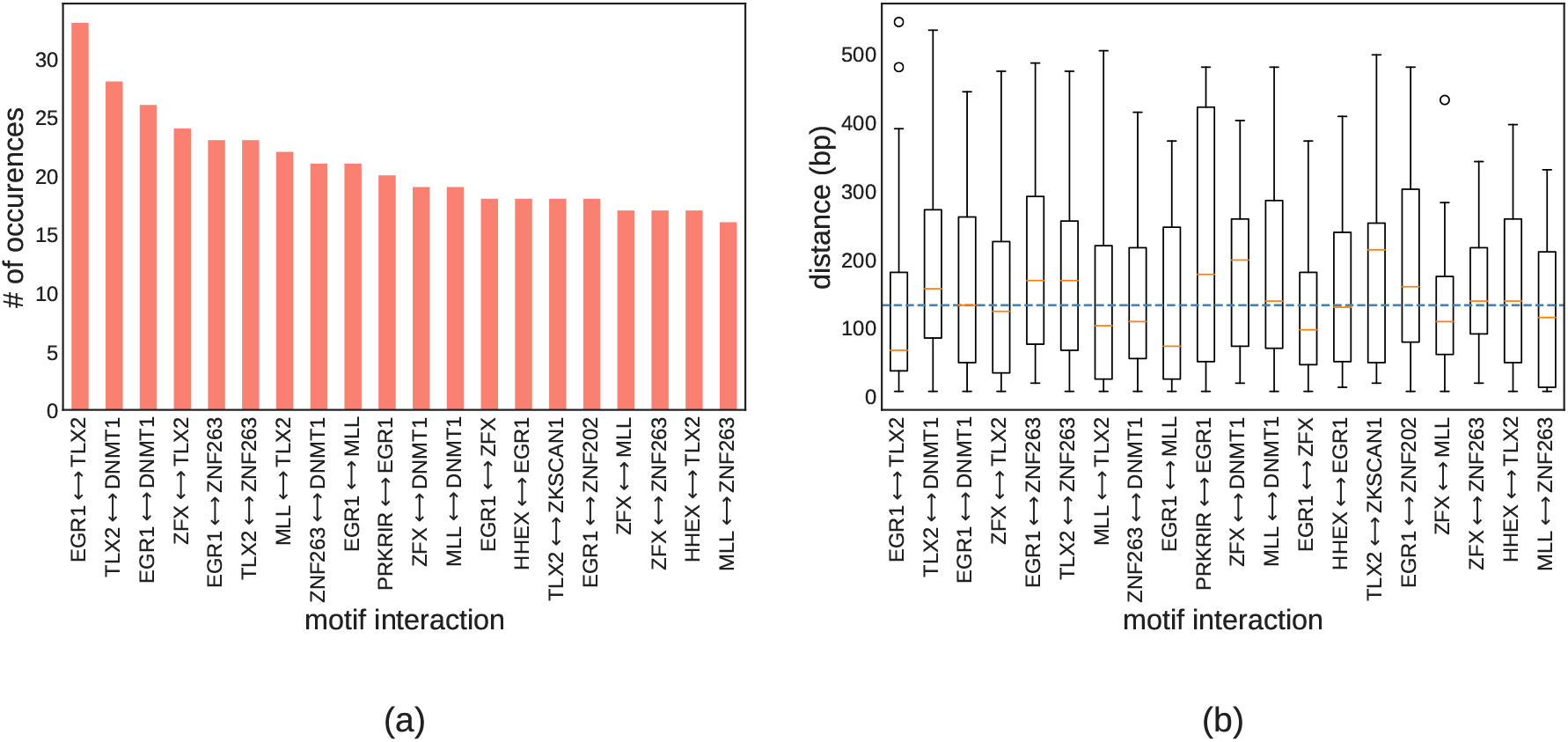
TF interactions. (a) The most frequent TF interactions in intron retention events. (b) The distribution of distances between detected TF interactions. The dotted blue line represents the median distance across all significant interactions.

## 3. Discussion

In our motif analysis we found that the C2H2 zinc finger family of TFs has a strong association with IR events: More than 50% of all motifs associated with IR have significant hits to C2H2 ZF TFs. This is consistent with previous work reporting that zinc finger TFs influence exon skipping [21]. This suggests that the C2H2 ZF family plays an important role in the regulation of alternative splicing in general.

To validate our predictions on the association of these TFs with IR, we used ChIP-seq data for multiple zinc finger TFs: MAZ, EGR1, SP1, ZBTB7A, SP2, and ZNF263.

We observed much higher occupancy of these TFs in IR events in the K562 human cell line, validating the model’s predictions. We also found that MAZ and EGR1 have specific preferences on where to bind within an IR/non-IR event: We observed that both TFs have stronger binding preference in the flanking exons of retained introns (see Figure 4). In contrast, in non-IR events, MAZ and EGR1 exhibit increased binding in the intronic region. This suggests a different mode of action for these TFs in IR vs non-IR events. Robson et al. [43] have reported that MAZ4 elements that contains four copies of the MAZ binding sequence influence alternative splicing [43]. More recently it was demonstrated that MAZ acts in conjunction with CTCF to remodel chromatin to affect changes in alternative splicing [44]. They have also demonstrated that like CTCF, MAZ can slow the elongation of RNAPII and affect splicing outcome.

There are multiple potential mechanisms by which TFs can affect co-transcriptional splicing. First, TFs are known to be critical in establishing chromatin state, which in turn can regulate alternative splicing by a purely kinetic model of the coupling between transcription and splicing whereby higher speeds of transcription in regions of accessible chromatin give less time for the spliceosomal machinery to recognize and splice those introns co-transcriptionally [11,45,46]. An alternative explanation of this phenomenon is that accessible chromatin is a mark of binding of TFs or other regulatory proteins that recruit splicing factors directly or indirectly through chromatin modifications to affect the outcome of splicing [7]. Wet-lab experiments are required to explore these hypotheses and provide more detailed information on how TFs affect intron retention, and splicing more generally, in a condition dependent manner.

Our model of retained introns considered only chromatin accessibility. There are other aspects of chromatin organization that can be considered: histone modifications and DNA methylation. Through their effect on chromatin organization, histone modifications impact the speed of RNAPII elongation and thereby alternative splicing [45]. Luco et al. [47] proposed the *adaptor system* model whereby DNA-binding proteins recognize a histone modification and recruit a splicing regulator that affects the splicing outcome (see also [48]). Methylation-dependent alternative splicing has been shown to be widespread [49], and its patterns have been observed to delineate exons and their boundaries [50,51]. Histone modifications and methylation patterns can thus provide another layer of information relevant to the regulation of IR.

In this work we focused on the local coupling of accessible chromatin and IR. We expect that non-local interactions through chromatin loop anchors like those that allow enhancers to affect promoter activity [52], can affect IR; evidence for their impact on exon skipping has recently been reported in human [53]. Recent work has demonstrated the role of a specific enhancer within a chromatin loop and its role in regulating alternative splicing [54]. Future work can incorporate them in the context of a comprehensive model of alternative splicing.

## 4. Conclusions and Future Work

Using deep learning to model intragenic DHSs allowed us to explore the regulatory elements that are predictive of IR in an unbiased fashion and identify TFs as key contributors to the regulation of IR. Further experimental work is required in order to validate the role of TFs in IR regulation. This will be supported by extensions of the model that allow condition-specific prediction of the IR state of regions of open chromatin, and create the chromatin-mediated IR code. Furthermore, the modularity of deep learning will allow the extension of the model to incorporate other sources of data indicative of chromatin state such as histone modifications. Much in the same way chromatin loop anchors allow enhancers to affect the activity of promoter regions and affect gene expression [52], there is recent evidence for their impact on exon skipping [53]. Therefore we expect that chromatin interaction information captured by Hi-C or Micro-C data is likely to improve the model and provide a more holistic view of IR regulation. Such data can be incorporated in a deep learning model with modules that use graph convolution; recent work has shown the effectiveness of this approach for modeling various aspects of chromatin state [55].

## 5. Methods

### 5.1. Data collection, processing, and representation

We use DNase I-seq data from 125 human immortalized cell-lines and tissues from the ENCODE database [56] and 39 cell types from the Roadmap Epigenetics consortium [57] as processed by [23]: every DNase I-seq peak is extended to a length of 600*bp* around its midpoint and adjacent peaks are greedily merged until no two peaks overlap by more than 200*bp*. For our analysis we focus on over a million DHSs that occur within genes.

Next, we extracted IR events from the Ensembl GRCh37 (hg19) reference annotations, utilizing code from SpliceGrapher [58] and iDiffIR [59]. In total, we identified 58, 305 unique IR events out of which, 15, 400 had overlapping DHSs. These constitute our positive examples. We use a strict criterion requiring the DHS to overlap the retained intron, i.e., DHSs overlapping only the flanking exons do not qualify. All other intragenic DHSs that did not overlap an IR event are labelled as negative examples. The number of negative examples was roughly twice the size of our positive set.

We use two methods to transform the sequences into input for the neural network: one-hot encoding and sequence embedding. For one-hot encoding a sequence is represented as a 4 × *N* matrix where *N* is the length of the sequence. Each position in the sequence is represented by the columns of the matrix with a non-zero value at a position corresponding to one of the four DNA nucleotides. To represent a sequence using embedding we first decompose it into overlapping *k*-mers of length *k*, and then use a word2vec model [60] to map each *k*-mer into an *m*-dimensional vector space. This gives us an embedding matrix of dimensions (*N − k* + 1) × *m*. This representation is designed to preserve the context of the *k*mers by producing similar embedding vectors for *k*-mers that tend to co-occur. Recently in a TF binding site prediction task within genomic sequences, it has been shown that in contrast to one-hot-encoding, k-mer embedding representation of the input leads to improved model performance [61].

### 5.2. Network architecture

We investigate several network architectures to predict chromatin accessibility in IR events with the goal of understanding its chromatin-mediated regulation. The primary network element, a one-dimensional convolutional layer, scans a set of filters against the matrix representing the input sequence. Formally, we can express the convolution operation as:

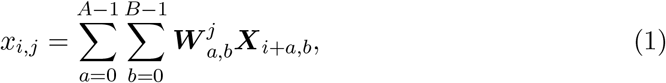

where ***X*** is the input matrix, *i* is the current output index, and *j* is the index of the filter. ***W*** is the weight matrix with size *A × B* where *A* is the length of the filter (window size) and *B* is the number of input channels: 4 for DNA one-hot encoding, *d* in case of word2vec embeddings, and *number of previous layer filters* in case of higher convolutional layers. The output of a convolutional layer is produced by applying a non-linear activation function to the result of the convolution operation. We use the Rectified Linear Unit (ReLU) which is given by:

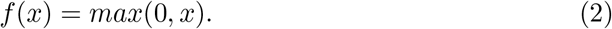

Next, the size of the output is reduced by max-pooling where the maximum value in a window of a pre-determined size is selected. This reduces the input size for the next layer and also leads to invariance to small shifts in the input sequence.

Another feature that we explore in our model are recurrent layers. RNNs have an internal state that enables them to capture distant feature interactions in the input sequence. Specifically, we employ a bi-directional RNN with Long Short-Term Memory (LSTM) units [62]. In a bi-directional RNN, a forward and a backward layer are used that traverse the input in both directions, improving the model’s performance.

We also incorporate a multi-head self-attention layer in our deep learning model. Attention is a powerful feature in that it can model dependencies within the input sequence regardless of their distances [63]. By doing so, it guides the network to focus on relevant features within the input and ignore irrelevant information. Our implementation uses code we have developed for the SATORI method [40].

### 5.3. Network training and evaluation

As mentioned in the previous section, we explore several network variants with different layers and features. The primary architecture used for the task at hand is shown Figure 2 (see supplementary materials for the summary of other architectures we explored). First the data is split into training, validation, and test sets with 80%, 10%, and 10% of the total data, respectively. Next, using the training and validation sets, we tune the network hyperparameters by employing a semi-randomized grid search algorithm that uses a 5-fold cross validation strategy. In case of the Basset-like model variant, we started with the hyperparaemters reported in [23] and fine-tuned their values. The optimized hyperparameters are summarized in supplementary table S1. To evaluate the model and the motif extraction analysis described later in this section, we use the test set. To assess model performance, we use the area under the ROC curve (AUC-ROC) and the area under the Precision-Recall curve (AUC-PRC).

### 5.4. Gapped kmer SVM

As a baseline we used the large-scale gapped kmer SVM (gkm-SVM), called the LS-GKM [37]. This version can handle bigger datasets (50k-100k examples) and exhibits better scalability. We run the package with the following parameters: − *m* 20000 and *T* − 16 which specify the size of the memory cache in MB and number of processing threads, respectively.

### 5.5. Motif extraction and analysis

To interpret the deep learning models, we extracted sequence motifs using the weights (filters) of the first convolutional layer, similar to the methodology described by Kelley et al. [23]. We select the positive examples (DHSs overlapping IR events) with model prediction probability greater than 0.65. This cutoff is chosen as a trade-off between the number of qualified examples and confidence in the prediction. For the negative examples, we used a cutoff value of less than 0.35. Next, for each filter we identify regions in the set of sequences that activated the filter with a value greater than half of the filter’s maximum score over all sequences. The highest scoring regions from all the sequences are stacked and for each filter, a position weight matrix is calculated using the nucleotide frequency and background information. We generate the sequence logos using the WebLogo tool [64]. The resulting PWMs are searched against the human CIS-BP database [38] using the TomTom tool [39] with distance metric set to Euclidean.

### 5.6. TF ChIP-seq analysis

We download the ChIP peaks of all the TFs that are enriched in IR events from the ENCODE database [56]. Next, we use our previously published pipeline [14] to test the enrichment of a given TF ChIP peaks in IR events. Briefly, we quantify the overlap of ChIP peaks with IR events and compare them to the overlap with non-IR events. The significance of overlap is tested using the Fisher exact test. To generate the profiles of TF occupancy across IR and non-IR events, we used the region of the ChIP peak where the PWM of the corresponding TF has the highest score. PWM scoring analysis was performed using BioPython [65].

### 5.7. Discovering interactions between TFs

To discover regulatory interactions between TFs we used SATORI [40], which takes advantage of the self-attention matrix to infer possible interactions between sequence motifs. When running SATORI, we used the default parameters with exception to the following: --attncutoff 0.08 and --usevalidtest True. The postprocesing was performed using Jupyter notebooks provided with SATORI.

## References

[1] A S N Reddy. Alternative splicing of pre-messenger RNAs in plants in the genomic era. Annu. Rev. Plant Biol., 58:267–94, 2007.

[2] A Kalsotra and TA Cooper. Functional consequences of developmentally regulated alternative splicing. Nature Rev. Genetics, 12:715–29, 2011.

[3] Geoffray Monteuuis, Justin JL Wong, Charles G Bailey, Ulf Schmitz, and John EJ Rasko. The changing paradigm of intron retention: regulation, ramifications and recipes. Nucleic acids research, 47(22):11497–11513, 2019.

[4] A S N Reddy, M F Rogers, D N Richardson, M Hamilton, and A Ben-Hur. Deciphering the plant splicing code: experimental and computational approaches for predicting alternative splicing and splicing regulatory elements. Front. Plant Sci., 3:18, 2012.

[5] Saurabh Chaudhary, Waqas Khokhar, Ibtissam Jabre, Anireddy SN Reddy, Lee J Byrne, Cornelia M Wilson, and Naeem H Syed. Alternative splicing and protein diversity: plants versus animals. Frontiers in plant science, 10:708, 2019.

[6] Justin J-L Wong, Amy YM Au, William Ritchie, and John EJ Rasko. Intron retention in mRNA: No longer nonsense. Bioessays, 38(1):41–49, 2016.

[7] U Braunschweig, N L Barbosa-Morais, Q Pan, E N Nachman, B Alipanahi, T Gonatopoulos-Pournatzis, B Frey, M Irimia, and B J Blencowe. Widespread intron retention in mammals functionally tunes transcription. Genome Research, 24:1774–86, 2014.

[8] Ying Ge and Bo T Porse. The functional consequences of intron retention: alternative splicing coupled to NMD as a regulator of gene expression. Bioessays, 36(3):236–243, 2014.

[9] Darya P Vanichkina, Ulf Schmitz, Justin J-L Wong, and John EJ Rasko. Challenges in defining the role of intron retention in normal biology and disease. In Seminars in Cell & Developmental Biology. Elsevier, 2017.

[10] Hyunchul Jung, Donghoon Lee, Jongkeun Lee, Donghyun Park, Yeon Jeong Kim, Woong-Yang Park, Dongwan Hong, Peter J Park, and Eunjung Lee. Intron retention is a widespread mechanism of tumor-suppressor inactivation. Nature genetics, 47(11):1242, 2015.

[11] Shiran Naftelberg, Ignacio E Schor, Gil Ast, and Alberto R Kornblihtt. Regulation of alternative splicing through coupling with transcription and chromatin structure. Annual Review of Biochemistry, 84:165–198, 2015.

[12] Fernando Carrillo Oesterreich, Lydia Herzel, Korinna Straube, Katja Hujer, Jonathon Howard, and Karla M Neugebauer. Splicing of nascent RNA coincides with intron exit from RNA polymerase II. Cell, 165(2):372–381, 2016.

[13] T R Mercer, S L Edwards, M B Clark, S J Neph, H Wang, A B Stergachis, S John, R Sandstrom, G Li, K S Sandhu, Y Ruan, L K Nielsen, J S Mattick, and J Stamatoyannopoulos. DNase I-hypersensitive exons colocalize with promoters and distal regulatory elements. Nature Genetics, 45:852–59, 2013.

[14] Fahad Ullah, Michael Hamilton, Anireddy SN Reddy, and Asa Ben-Hur. Exploring the relationship between intron retention and chromatin accessibility in plants. BMC genomics, 19(1):21, 2018.

[15] R E Thurman, E Rynes, R Humbert, J Vierstra, M T Maurano, E Haugen, N C Sheffield, A B Stergachis, H Wang, B Vernot, K Garg, S John, R Sandstrom, D Bates, L Boatman, T K Canfield, M Diegel, D Dunn, A K Ebersol, T Frum, E Giste, A K Johnson, E M Johnson, T Kutyavin, B Lajoie, B K Lee, K Lee, D London, D Lotakis, S Neph, F Neri, E D Nguyen, H Qu, A P Reynolds, V Roach, A Safi, M E Sanchez, A Sanyal, A Shafer, J M Simon, L Song, S Vong, M Weaver, Y Yan, Z Zhang, B Lenhard, M Tewari, M O Dorschner, R S Hansen, P A Navas, G Stamatoyannopoulos, V R Iyer, J D Lieb, S R Sunyaev, J M Akey, P J Sabo, R Kaul, T S Furey, J Dekker, G E Crawford, and J A Stamatoyannopoulos. The accessible chromatin landscape of the human genome. Nature, 489(7414):75–82, 2012.

[16] G Felsenfeld and M Groudine. Controlling the double helix. Nature, 421:448–53, 2003.

[17] D S Gross and W T Garrard. Nuclease hypersensitive sites in chromatin. Annual Rev. Biochem., 57:159–197, 1988.

[18] D J Galas and A Schmitz. DNase footprinting: A simple method for detection of protein-DNA binding specificity. Nucleic Acids Res., 5:3157–70, 1978.

[19] J R Hesselberth, X Y Chen, Z H Zhang, P J Sabo, R Sandstrom, A P Reynolds, R E Thurman, S Neph, M S Kuehn, W S Noble, S Fields, and J A Stamatoyannopoulos. Global mapping of protein-DNA interactions in-vivo by digital genomic footprinting. Nature Methods, 6:283–89, 2009.

[20] A P Boyle, L Y Song, B K Lee, D London, D Keefe, E Birney, V R Iyer, G E Crawford, and T S Furey. High-resolution genome-wide in-vivo footprinting of diverse transcription factors in human cells. Genome Research, 21:456–64, 2011.

[21] Hong Han, Ulrich Braunschweig, Thomas Gonatopoulos-Pournatzis, Robert J Weatheritt, Calley L Hirsch, Kevin CH Ha, Ernest Radovani, Syed Nabeel-Shah, Tim Sterne-Weiler, Juli Wang, et al. Multilayered control of alternative splicing regulatory networks by transcription factors. Molecular cell, 65(3):539–553, 2017.

[22] Steven J Burgess, Ivan Reyna-Llorens, Sean R Stevenson, Pallavi Singh, Katja Jaeger, and Julian M Hibberd. Genome-wide transcription factor binding in leaves from C3 and C4 grasses. The Plant Cell, 31(10):2297–2314, 2019.

[23] David R Kelley, Jasper Snoek, and John L Rinn. Basset: learning the regulatory code of the accessible genome with deep convolutional neural networks. Genome research, 26(7):990–999, 2016.

[24] Nicholas E Banovich, Yang I Li, Anil Raj, Michelle C Ward, Peyton Greenside, Diego Calderon, Po Yuan Tung, Jonathan E Burnett, Marsha Myrthil, Samantha M Thomas, et al. Impact of regulatory variation across human iPSCs and differentiated cells. Genome research, 28(1):122–131, 2018.

[25] Christof Angermueller, Heather J Lee, Wolf Reik, and Oliver Stegle. DeepCpG: accurate prediction of single-cell DNA methylation states using deep learning. Genome biology, 18(1):67, 2017.

[26] Zeynep Kalender Atak, Ibrahim Ihsan Taskiran, Jonas Demeulemeester, Christopher Flerin, David Mauduit, Liesbeth Minnoye, Gert Hulselmans, Valerie Christiaens, Ghanem-Elias Ghanem, Jasper Wouters, and Stein Aerts. Interpretation of allele-specific chromatin accessibility using cell state–aware deep learning. Genome research, 31(6):1082–1096, 2021.

[27] Babak Alipanahi, Andrew Delong, Matthew T Weirauch, and Brendan J Frey. Predicting the sequence specificities of DNA-and RNA-binding proteins by deep learning. Nature biotechnology, 33(8):831, 2015.

[28] Qian Qin and Jianxing Feng. Imputation for transcription factor binding predictions based on deep learning. PLoS computational biology, 13(2):e1005403, 2017.

[29] Ameni Trabelsi, Mohamed Chaabane, and Asa Ben-Hur. Comprehensive evaluation of deep learning architectures for prediction of DNA/RNA sequence binding specificities. Bioinformatics, 35(14):i269–i277, 2019.

[30] Pang Wei Koh, Emma Pierson, and Anshul Kundaje. Denoising genome-wide histone chip-seq with convolutional neural networks. Bioinformatics, 33(14):i225–i233, 2017.

[31] Jacob Schreiber, Maxwell Libbrecht, Jeffrey Bilmes, and William Noble. Nucleotide sequence and DNaseI sensitivity are predictive of 3D chromatin architecture. bioRxiv, page 103614, 2018.

[32] Xiaoyong Pan and Hong-Bin Shen. RNA-protein binding motifs mining with a new hybrid deep learning based cross-domain knowledge integration approach. BMC bioinformatics, 18(1):136, 2017.

[33] Sai Zhang, Jingtian Zhou, Hailin Hu, Haipeng Gong, Ligong Chen, Chao Cheng, and Jianyang Zeng. A deep learning framework for modeling structural features of RNA-binding protein targets. Nucleic acids research, 44(4):e32–e32, 2015.

[34] Michael KK Leung, Hui Yuan Xiong, Leo J Lee, and Brendan J Frey. Deep learning of the tissue-regulated splicing code. Bioinformatics, 30(12):i121–i129, 2014.

[35] Anupama Jha, Matthew R Gazzara, and Yoseph Barash. Integrative deep models for alternative splicing. Bioinformatics, 33(14):i274–i282, 2017.

[36] Kishore Jaganathan, Sofia Kyriazopoulou Panagiotopoulou, Jeremy F McRae, Siavash Fazel Darbandi, David Knowles, Yang I Li, Jack A Kosmicki, Juan Arbelaez, Wenwu Cui, Grace B Schwartz, et al. Predicting splicing from primary sequence with deep learning. Cell, 2019.

[37] Dongwon Lee. LS-GKM: a new gkm-svm for large-scale datasets. Bioinformatics, 32 14:2196–8, 2016.

[38] Matthew T. Weirauch, Ally Yang, Mihai Albu, Atina G. Cote, Alejandro Montenegro-Montero, Philipp Drewe, Hamed S. Najafabadi, Samuel A. Lambert, Ishminder Mann, Kate B. Cook, Hong Yuan Zheng, Alejandra Goity, Harm van Bakel, Javier Fernández Lozano, Mary Galli, Mathew G Lewsey, Eryong Huang, Tuhin Mukherjee, Xiaoting Chen, John S. Reece-Hoyes, Sridhar Govindarajan, Gad Shaulsky, Albertha J M Walhout, François-Yves Bouget, Gunnar Ratsch, Luis F Larrondo, Joseph R. Ecker, and Timothy R. Hughes. Determination and inference of eukaryotic transcription factor sequence specificity. Cell, 158:1431–1443, 2014.

[39] Shobhit Gupta, John A. Stamatoyannopoulos, Timothy L. Bailey, and William Stafford Noble. Quantifying similarity between motifs. Genome Biology, 8:R24–R24, 2006.

[40] Fahad Ullah and Asa Ben-Hur. A self-attention model for inferring cooperativity between regulatory features. Nucleic Acids Research, 49(13):e77, 2021.

[41] Heonjong Han, Jae-Won Cho, Sangyoung Lee, Ayoung Yun, Hyojin Kim, Dasom Bae, Sunmo Yang, Chan Yeong Kim, Muyoung Lee, Eunbeen Kim, et al. TRRUST v2: an expanded reference database of human and mouse transcriptional regulatory interactions. Nucleic acids research, 46(D1):D380–D386, 2018.

[42] Gregorio Alanis-Lobato, Miguel A Andrade-Navarro, and Martin H Schaefer. HIPPIE v2.0: enhancing meaningfulness and reliability of protein–protein interaction networks. Nucleic acids research, page gkw985, 2016.

[43] Nicole D Robson-Dixon and Mariano A Garcia-Blanco. MAZ elements alter transcription elongation and silencing of the fibroblast growth factor receptor 2 exon IIIb. Journal of Biological Chemistry, 279(28):29075–29084, 2004.

[44] Tiaojiang Xiao, Xin Li, and Gary Felsenfeld. The Myc-associated zinc finger protein (MAZ) works together with CTCF to control cohesin positioning and genome organization. Proceedings of the National Academy of Sciences, 118(7), 2021.

[45] Tassa Saldi, Michael A Cortazar, Ryan M Sheridan, and David L Bentley. Coupling of RNA polymerase II transcription elongation with pre-mRNA splicing. Journal of molecular biology, 428(12):2623–2635, 2016.

[46] Heidi Dvinge. Regulation of alternative mRNA splicing: old players and new perspectives. FEBS letters, 2018.

[47] Reini F Luco, Qun Pan, Kaoru Tominaga, Benjamin J Blencowe, Olivia M Pereira-Smith, and Tom Misteli. Regulation of alternative splicing by histone modifications. Science, 327(5968):996–1000, 2010.

[48] I E Schor, M Allo, and A R Kornblihtt. Intragenic chromatin modifications: A new layer in alternative splicing regulation. Epigenetics, 5(3):174–179, 2010.

[49] J Wan, V F Oliver, H Zhu, D J Zack, J Qian, and S L Merbs. Integrative analysis of tissue-specific methylation and alternative splicing identifies conserved transcription factor binding motifs. Nucleic Acids Res, 41(18):8503–8514, 2013.

[50] Sahar Gelfman, Noa Cohen, Ahuvi Yearim, and Gil Ast. DNA-methylation effect on cotranscriptional splicing is dependent on GC architecture of the exon–intron structure. Genome research, 23(5):789–799, 2013.

[51] Galit Lev Maor, Ahuvi Yearim, and Gil Ast. The alternative role of DNA methylation in splicing regulation. Trends in Genetics, pages 1–7, 2015.

[52] William W Greenwald, He Li, Paola Benaglio, David Jakubosky, Hiroko Matsui, Anthony Schmitt, Siddarth Selvaraj, Matteo D’Antonio, Agnieszka D’Antonio-Chronowska, Erin N Smith, and Kelly A Frazer. Subtle changes in chromatin loop contact propensity are associated with differential gene regulation and expression. Nature communications, 10(1):1–17, 2019.

[53] Yu Zhang, Yichao Cai, Xavier Roca, Chee Keong Kwoh, and Melissa Jane Fullwood. Chromatin loop anchors predict transcript and exon usage. Briefings in Bioinformatics, 2021.

[54] Sara Dahan, Aveksha Sharma, Klil Cohen, Mai Baker, Nadeen Taqatqa, Mercedes Bentata, Eden Engal, Ahmad Siam, Gillian Kay, Yotam Drier, Shlomo Elias, and Maayan Salton. VEGFA’s distal enhancer regulates its alternative splicing in CML. NAR cancer, 3(3):zcab029, 2021.

[55] Jack Lanchantin and Yanjun Qi. Graph convolutional networks for epigenetic state prediction using both sequence and 3D genome data. Bioinformatics, 36:i659–i667, 2020.

[56] Avinash Das Sahu. An integrated encyclopedia of DNA elements in the human genome. 2012.

[57] Anshul Kundaje, Wouter Meuleman, Jason Ernst, Misha Bilenky, Angela Yen, Alireza Heravi-Moussavi, Pouya Kheradpour, Zhizhuo Zhang, Jianrong Wang, Michael J Ziller, et al. Integrative analysis of 111 reference human epigenomes. In Nature, 2015.

[58] M F Rogers, J Thomas, A S N Reddy, and A Ben-Hur. SpliceGrapher: Detecting patterns of alternative splicing from RNA-seq data in the context of gene models and EST data. Genome Biology, 13:1–17, 2012.

[59] Sergei A Filichkin, Michael Hamilton, Palitha D Dharmawardhana, Sunil K Singh, Christopher Sullivan, Asa Ben-Hur, Anireddy SN Reddy, and Pankaj Jaiswal. Abiotic stresses modulate landscape of poplar transcriptome via alternative splicing, differential intron retention, and isoform ratio switching. Frontiers in plant science, 9:5, 2018.

[60] Tomas Mikolov, Ilya Sutskever, Kai Chen, Greg S Corrado, and Jeff Dean. Distributed representations of words and phrases and their compositionality. In Advances in neural information processing systems, pages 3111–3119, 2013.

[61] Zhen Shen, Wenzheng Bao, and De-Shuang Huang. Recurrent neural network for predicting transcription factor binding sites. Scientific reports, 8(1):1–10, 2018.

[62] Sepp Hochreiter and Jörgen Schmidhuber. Long short-term memory. Neural computation, 9(8):1735–1780, 1997.

[63] Ashish Vaswani, Noam Shazeer, Niki Parmar, Jakob Uszkoreit, Llion Jones, Aidan N Gomez, Łukasz Kaiser, and Illia Polosukhin. Attention is all you need. In Advances in neural information processing systems, pages 5998–6008, 2017.

[64] G E Crooks, G Hon, J M Chandonia, and S E Brenner. Weblogo: A sequence logo generator. Genome Research, 14:1188–90, 2004.

[65] Peter JA Cock, Tiago Antao, Jeffrey T Chang, Brad A Chapman, Cymon J Cox, Andrew Dalke, Iddo Friedberg, Thomas Hamelryck, Frank Kauff, Bartek Wilczynski, et al. Biopython: freely available python tools for computational molecular biology and bioinformatics. Bioinformatics, 25(11):1422–1423, 2009.

