## Supplement for "Evidence for the role of transcription factors in the co-transcriptional regulation of intron retention"

#### Generating transcription factor family distributions across the human genome

To generate Figure 1 in the main text, we used the Basset network architecture [1] with all DHSs (~2 million) across 164 human cell lines. The dataset was split into 80%, 10%, and 10% for training, validating, and testing. Once the model was trained, we followed the motif analysis pipeline described in [1]. We analyzed first CNN layer filters to generate motifs for three different test sets: DHSs that overlapped human promoters, intragenic, and intergenic regions, separately. Essentially, for each genomic region, we identified regions in the corresponding set of sequences that activated the filter with a value greater than half of the filter's maximum score over all sequences. The highest scoring regions from all the sequences were stacked and for each filter, a position weight matrix was calculated using the nucleotide frequency and background information. The motifs learned by the network were mapped to the human CISBP database [2] using the TomTom tool from the MEME suite [3] with distance metric set to Euclidean. Finally, we picked the significantly matched transcription factors (adjusted p-value $< 0.05$) and annotated them according to their families.

#### Embeddings and network interpretability

We used the Basset-like network (see Figure 2 in the main text) to demonstrate that embeddings reduce network interpretability. We used this model with both one-hot and word2vec representations of the input sequences. Interestingly, the average information content (IF) of enriched motifs significantly varied with the two input representations. When using the regular one-hot encoding, we find the motifs to be more informative and useful (mean IF = 4.0). The same is not true for word2vec embeddings where we get motifs with far lower information content (mean IF = 1.8).

### Supplementary Figures

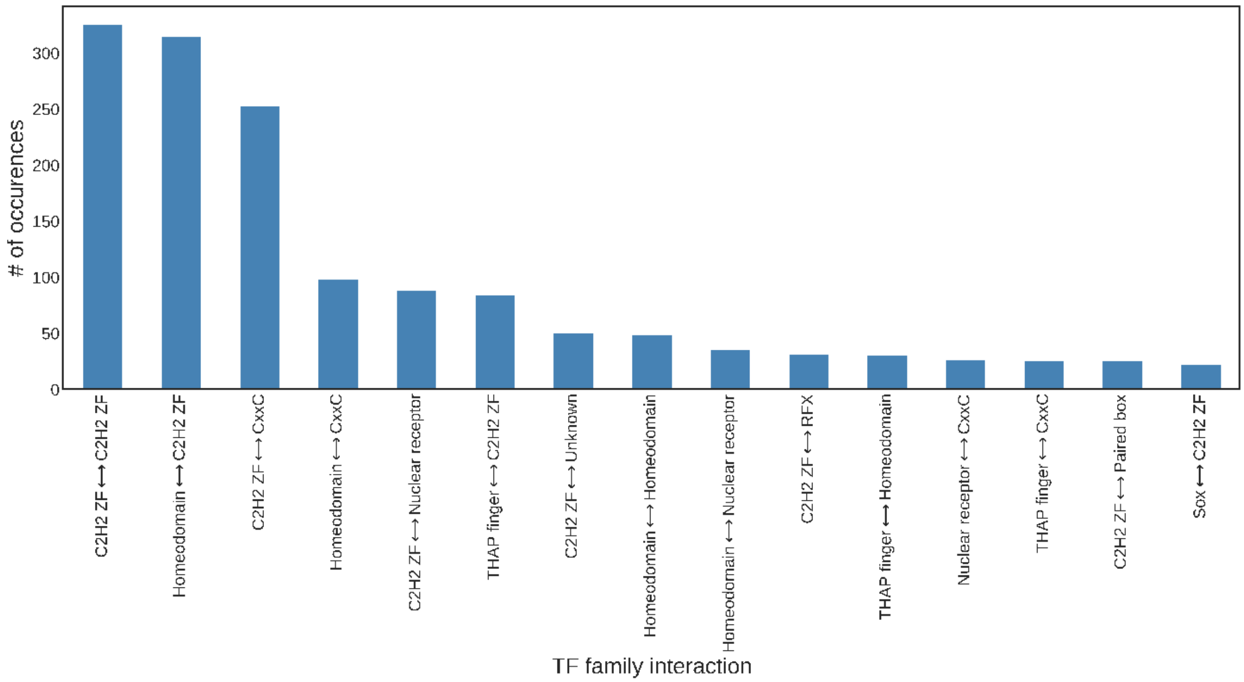


**Figure F1:** The most frequent interacting transcription factor families in the intron retention events.

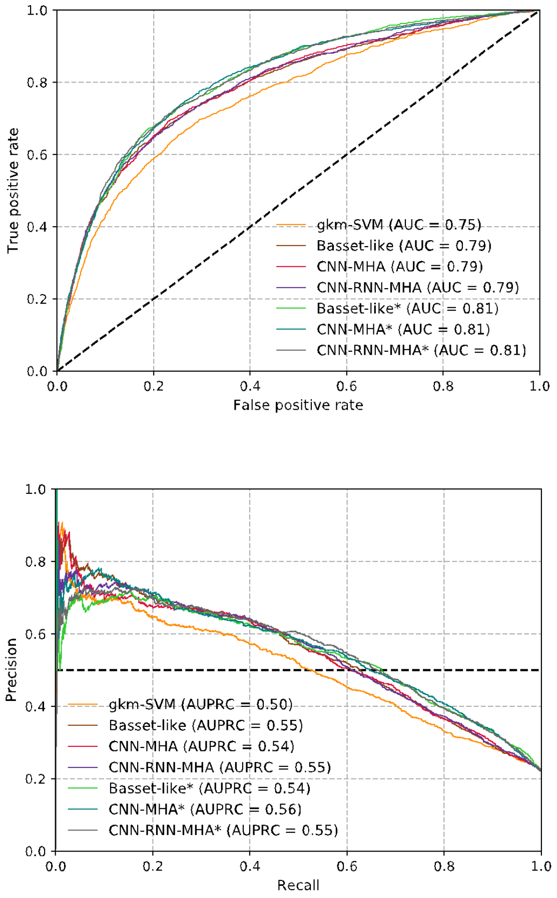


**Figure F2:** ROC (left) and Precision-Recall (right) curves for the different deep learning architectures as well as the gkm-SVM. Network architectures that use *k*-mer embeddings instead of one-hot encoding are indicated by an asterisk (*). The AUC and AUPRC values are provided in the legends.

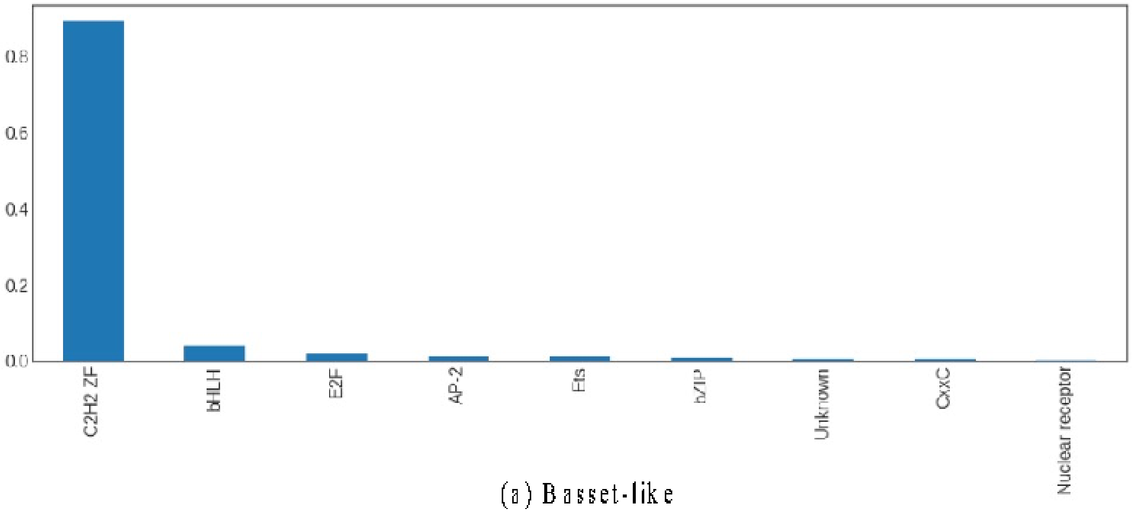


(a) Basset-like

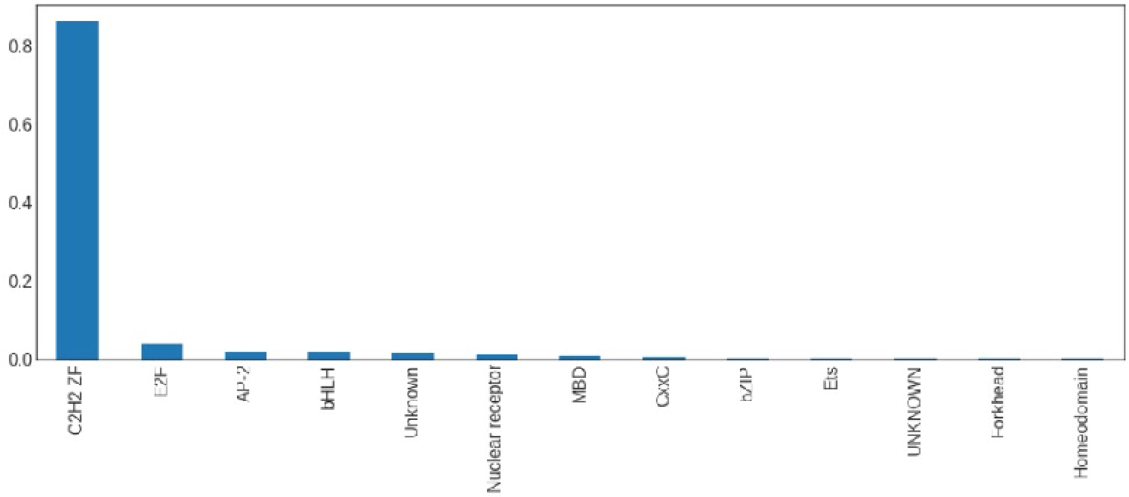


(b) CNN-MHA

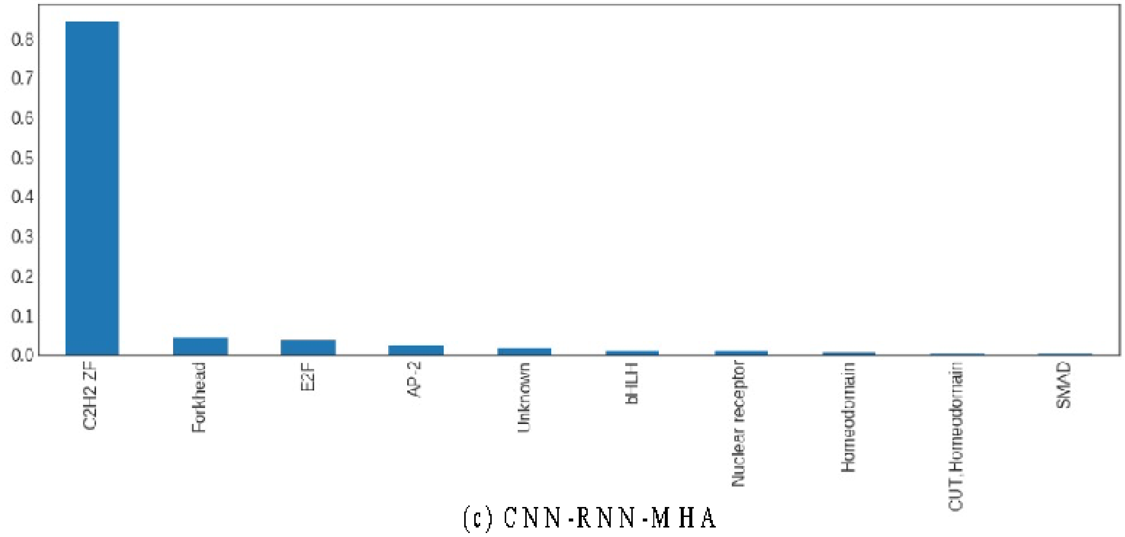


(c) CNN-RNN-MHA

**Figure F4:** Families of motifs identified in IR events are shown for (a) Basset-like, (b) CNN-MHA, and (c) CNN-RNN-MHA architectures, sorted by the percentage of all identified motif families. It follows that regardless of the architecture, C2H2 ZF is the most frequent family of transcription factors enriched in IR events.

#### Supplementary Tables

**Table S1:** List of network hyperparameters.

| Hyperparameter | Type | Description |
|---|---|---|
| singlehead_size | int | Size of the attention single head [default: 32] |
| num_heads | int | Number of heads in multi-head self-attention layer [default: 8] |
| multihead_size | int | Output size of the multi-head after concatenation [default: 100] |
| batch_size | int | Batch size in training/testing the model [default: 172] |
| use_RNN | bool | Choose whether to use the RNN layer. [default: based on model variant] |
| RNN_hidden_size | int | Size of the RNN layer. [default: 100] |
| CNN_filters | int | Number of CNN filters to use. [default: 200] |
| CNN_filter_size | int | Size of each CNN filter. [default: 13] |
| use_CNN_pool | bool | Use max pooling in the CNN layer. [default: True] |
| CNN_pool_size | int | Size of the max pooling window in CNN layer. [default: 6] |
| input_channels | int | Number of input channels. [default: 4 (for DNA sequences)] |
| num_epochs | int | Number of training epochs. [default: 30] |
| readout_strategy | string | Either to normalize the MHA output or flatten it. [default: "normalize"] |
| use_embd | bool | Whether to use the word2vec embeddings instead of 1-hot input [default: False] |
| embd_size | int | Size of the word2vec embedding vectors [default: 50] |
| embd_window | int | Size of the word2vec embedding window [default: 5] |
| embd_kmer | int | Length of the *k*mer (for word2vec embeddings) [default: 3] |

**Table S2:** A list of all significant interactions in intron retention events.
(provided in the additional Excel file)

**Table S3:** A list of TF interactions in IR events confirmed by the TRRUSTv2 database [4]. The level of significance(adjusted p-value) assigned by SATORI to each interaction is provided in the adusted_pval column.

| TF_Interaction | TF1_Family | TF2_Family | adjusted_pval | frequency |
|---|---|---|---|---|
| DNMT1↔HINFP | CxxC | C2H2 ZF | 2.39E-49 | 5 |
| EGR1↔DNMT1 | C2H2 ZF | CxxC | 3.68E-45 | 26 |
| ESR1↔DNMT1 | Nuclear receptor | CxxC | 5.58E-45 | 5 |
| PAX8↔DNMT1 | Paired box | CxxC | 4.63E-42 | 4 |
| ESR1↔PURA | Nuclear receptor | Unknown | 5.14E-36 | 2 |
| DNMT1↔ZBTB7A | CxxC | C2H2 ZF | 1.49E-33 | 5 |
| EGR1↔THRB | C2H2 ZF | Nuclear receptor | 7.77E-33 | 9 |
| DNMT1↔SP4 | C2H2 ZF | CxxC | 5.54E-30 | 4 |
| ESR1↔PAX8 | Nuclear receptor | Paired box | 1.48E-27 | 1 |
| HHEX↔PAX8 | Paired box | Homeodomain | 3.69E-22 | 4 |
| E2F4↔DNMT1 | E2F | CxxC | 1.29E-19 | 1 |
| ESR1↔EGR1 | Nuclear receptor | C2H2 ZF | 8.77E-15 | 6 |
| ESR1↔RARG | Nuclear receptor | Nuclear receptor | 1.03E-14 | 1 |
| ESR1↔E2F4 | Nuclear receptor | E2F | 1.61E-14 | 1 |
| EGR1↔PAX8 | Paired box | C2H2 ZF | 5.77E-13 | 5 |
| EGR1↔PURA | Unknown | C2H2 ZF | 8.00E-13 | 9 |
| ESR1↔ZBTB7A | Nuclear receptor | C2H2 ZF | 3.01E-07 | 1 |
| ESR1↔SP4 | Nuclear receptor | C2H2 ZF | 1.26E-06 | 1 |
| PURA↔SP4 | Unknown | C2H2 ZF | 3.50E-06 | 1 |
| EGR1↔E2F4 | E2F | C2H2 ZF | 1.08E-05 | 2 |
| EGR1↔SP4 | C2H2 ZF | C2H2 ZF | 2.36E-03 | 3 |

**Table S4:** A list of TF interactions in the IR events confirmed by the HIPPIE database [5]. The level of significance (adjusted p-value) assigned by SATORI to each interaction is provided in adjusted_pval column.

| TF_Interaction | TF1_Family | TF2_Family | adjusted_pval | frequency |
|---|---|---|---|---|
| ESR1⟷IRX4 | Nuclear receptor | Homeodomain | 5.95E-65 | 1 |
| ESR1⟷DNMT1 | Nuclear receptor | CxxC | 5.58E-45 | 5 |
| HHEX⟷TLX2 | Homeodomain | Homeodomain | 4.43E-36 | 17 |
| ESR1⟷PURA | Nuclear receptor | Unknown | 5.14E-36 | 2 |
| ESR1⟷ZHX1 | Nuclear receptor | Homeodomain | 8.48E-31 | 1 |
| SOX1⟷ESR1 | Sox | Nuclear receptor | 3.88E-28 | 2 |
| ESR1⟷PAX8 | Nuclear receptor | Paired box | 1.48E-27 | 1 |
| ESR1⟷HINFP | Nuclear receptor | C2H2 ZF | 1.41E-23 | 1 |
| ESR1⟷SP7 | Nuclear receptor | C2H2 ZF | 3.41E-17 | 3 |
| ESR1⟷EGR1 | Nuclear receptor | C2H2 ZF | 8.77E-15 | 6 |
| ESR1⟷RARG | Nuclear receptor | Nuclear receptor | 1.03E-14 | 1 |
| ESR1⟷E2F4 | Nuclear receptor | E2F | 1.61E-14 | 1 |
| EGR1⟷HINFP | C2H2 ZF | C2H2 ZF | 1.84E-14 | 8 |
| EGR1⟷PURA | Unknown | C2H2 ZF | 8.00E-13 | 9 |
| ZHX1⟷HINFP | Homeodomain | C2H2 ZF | 2.93E-09 | 1 |
| E2F4⟷HINFP | E2F | C2H2 ZF | 1.18E-08 | 1 |
| PURA⟷SP4 | Unknown | C2H2 ZF | 3.50E-06 | 1 |

**Table S5:** Mean and median information content for IR and non-IR filters are provided for the three architectures; Basset-like, convolution and multi-head attention (CNN-MHA), and convolution with both recurrent network and multi-head attention (CNN-RNN-MHA).

| Architecture | IR filters | | Non-IR filters | |
|---|---|---|---|---|
| | Mean | Median | Mean | Median |
| Basset-like | 4.12 | 4.21 | 4.26 | 4.26 |
| CNN-MHA | 4 | 4 | 4.4 | 4.38 |
| CNN-RNN-MHA | 3.9 | 3.85 | 4.42 | 4.41 |

### References

1. Kelley, D. R., Snoek, J., & Rinn, J. L. (2016). Basset: learning the regulatory code of the accessible genome with deep convolutional neural networks. *Genome research*, *26*(7), 990-999.
2. Lambert, S. A., Jolma, A., Campitelli, L. F., Das, P. K., Yin, Y., Albu, M., ... & Weirauch, M. T. (2018). The human transcription factors. *Cell*, *172*(4), 650-665.
3. Bailey, T. L., Johnson, J., Grant, C. E., & Noble, W. S. (2015). The MEME suite. *Nucleic acids research*, *43*(W1), W39-W49.
4. Han, H., Cho, J. W., Lee, S., Yun, A., Kim, H., Bae, D., ... & Lee, S. (2018). TRRUST v2: an expanded reference database of human and mouse transcriptional regulatory interactions. *Nucleic acids research*, *46*(D1), D380-D386.
5. Alanis-Lobato, G., Andrade-Navarro, M. A., & Schaefer, M. H. (2016). HIPPIE v2. 0: enhancing meaningfulness and reliability of protein–protein interaction networks. *Nucleic acids research*, gkw985.
